# Transient opening of tricellular vertices controls paracellular transport through the follicle epithelium during *Drosophila* oogenesis

**DOI:** 10.1101/2020.02.29.971168

**Authors:** Jone Isasti-Sanchez, Fenja Münz-Zeise, Stefan Luschnig

## Abstract

Paracellular permeability is regulated to allow solute transport or migration of cells across epithelial or endothelial barriers. However, how occluding junction dynamics controls paracellular permeability is poorly understood. Here we describe patency, a developmentally regulated process in *Drosophila* oogenesis, during which cell vertices in the follicle epithelium open transiently to allow paracellular transport of yolk proteins for uptake by the oocyte. We show that the sequential removal of E-Cadherin, N-Cadherin, NCAM/Fasciclin-2 and Sidekick from vertices precedes their basal-to-apical opening, while the subsequent assembly of tricellular occluding junctions terminates patency and seals the paracellular barrier. E-Cadherin-based adhesion is required to limit paracellular channel size, whereas stabilized adherens junctions, prolonged NCAM/Fasciclin-2 expression, impeded endocytosis, or increased actomyosin contractility prevent patency. Our findings reveal a key role of cell vertices as gateways controlling paracellular transport, and demonstrate that the dynamic regulation of adhesion and actomyosin contractility at vertices governs epithelial barrier properties.

## Introduction

Epithelia are selectively permeable tissue barriers that protect the body from the external environment, *e.g*. by lining the skin or the intestine, and they control the transport of substances between different body compartments to maintain tissue homeostasis. Ions and non-charged solutes move across epithelia either through cells (transcellular route) or through the space between cells (paracellular route). The transcellular pathway involves energy-dependent transport across the plasma membrane through pumps or carriers, often coupled with passive diffusion through ion channels, whereas substances that cross epithelia via the paracellular pathway diffuse along their electro-chemical gradient between the cells. Transport along this pathway is passive and is regulated by occluding junctions that connect adjacent cells and seal the space between them (Van Itallie and Anderson, 2006). Tight junctions (TJs; Zihni et al., 2016) in vertebrates and septate junctions (SJs; Izumi and Furuse, 2014) in invertebrates play functionally equivalent roles as occluding junctions in barrier epithelia. They restrict the passage of molecules through the paracellular space and separate apical and basolateral membrane domains in epithelial cells. While bicellular SJs and TJs seal contacts between two adjacent cells, specialized tricellular junctions (TCJs) form at cell vertices, where a distinct set of proteins mediates adhesive and occluding properties and controls paracellular permeability (Higashi and Miller, 2017; Bosveld et al., 2018). In *Drosophila*, the transmembrane proteins Anakonda (Aka; Byri et al., 2015) and Gliotactin (Gli; Schulte et al., 2003) are components of tricellular occluding junctions essential for barrier formation, while the adhesion protein Sidekick (Sdk; Lye et al., 2014) localizes specifically to vertices at the level of adherens junctions (AJs).

Cellular junctions are remodeled to mediate specific transport functions or to allow the passage of migrating cells. For instance, paracellular extravasation of leukocytes from blood vessels requires the transient opening of endothelial junctions stimulated by pro-inflammatory cytokines (Vestweber, 2015). In the testis, TJs between Sertoli cells constituting the blood-testis barrier (BTB) open transiently to allow translocation of developing spermatocytes across the seminiferous epithelium (Smith and Braun, 2012), involving the remodeling of TJ components controlled by androgens (Chakraborty et al., 2014). Moreover, certain bacterial pathogens, including Group A *Streptococcus*, breach paracellular barriers via TCJs, employing cleavage of TJ proteins (Sumitomo et al., 2016). Collectively, these processes require changes to the composition and dynamics of occluding junctions, but the underlying cellular and molecular mechanisms that mediate the opening and closing of junctions are not well understood.

Junctional remodeling is also involved in epithelial morphogenesis during development. Here we investigated the regulation of barrier function in the follicle cell epithelium (FCE) during oogenesis in *Drosophila*. Fly ovaries contain several ovarioles with successively maturing follicles (egg chambers), each of which comprises 16 germline cells (15 nurse cells and one oocyte) covered by an epithelial layer of somatic follicle cells (FCs; Duhart et al., 2017). The FCs proliferate until stage 6, when the activation of Notch signaling in the FCs triggers a cell cycle switch from mitotic to endoreplication cycles (Deng et al., 2001; Lopez-Schier and St Johnston, 2001). At mid-oogenesis the majority of FCs form a columnar epithelium covering the oocyte located in the posterior part of the egg chamber, while the FCs remaining in the anterior form a squamous epithelium covering the nurse cells (Duhart et al., 2017). At the same time the oocyte begins to increase in volume as a result of the uptake of cytoplasm from the nurse cells, which subsequently degenerate, and of yolk, which fills most of the oocyte volume. Yolk uptake (vitellogenesis) begins at stage 8, marking a key point during oogenesis associated with dramatic changes in FC morphology and physiology (Duhart et al., 2017). In higher dipterans, yolk consists of three major yolk proteins (YP1-3), which associate with lipids to form lipoprotein particles that provide energy reserves for the developing embryo (Raikhel and Dhadialla, 1992). However, unlike the cytoplasmic components of the oocyte, the yolk proteins are not synthesized by the nurse cells, but predominantly by the fat body, the equivalent of the vertebrate liver and adipose tissue, and to a minor extent by the FCs (Brennan et al., 1982). As the bulk of yolk protein is secreted by the fat body into the hemolymph, these proteins must pass the FCE before they can be taken up by the oocyte through receptor-mediated endocytosis (Raikhel and Dhadialla, 1992). Transport of yolk proteins from the hemolymph across the FCE is thought to occur via the paracellular route, enabled by the transient opening of channels between FCs, a condition referred to as *patency* (Patchin and Davey, 1968; Pratt and Davey, 1972). However, the cellular and molecular mechanisms underlying patency are not clear.

Here we investigated the cellular events and underlying mechanisms that lead to the transient opening of the FCE paracellular barrier during vitellogenesis. We show that the successive removal of adhesion proteins from cell vertices precedes the opening of intercellular spaces between FCs and that the subsequent formation of TCJs marks the termination of patency. Our findings reveal a key role of cell vertices in the regulation of paracellular transport across an epithelial barrier.

## Results

### Transient opening of cell vertices enables paracellular transport through the follicle epithelium during vitellogenesis

To investigate barrier function of the follicle epithelium during oogenesis we incubated living egg chambers in medium containing fluorescent dextran (10 or 40 kDa) as a tracer and the lipid dye MM4-64 to label cell membranes. We focused on the mainbody FCs, which are cuboidal in pre-vitellogenic follicles (stage 1-7), become columnar during vitellogenesis (stages 8-10B), and flatten during late vitellogenesis (stage 11; Fig. 1A-C). As the FCE undergoes these changes, the oocyte increases dramatically in volume due to yolk uptake.

**Figure 1.**
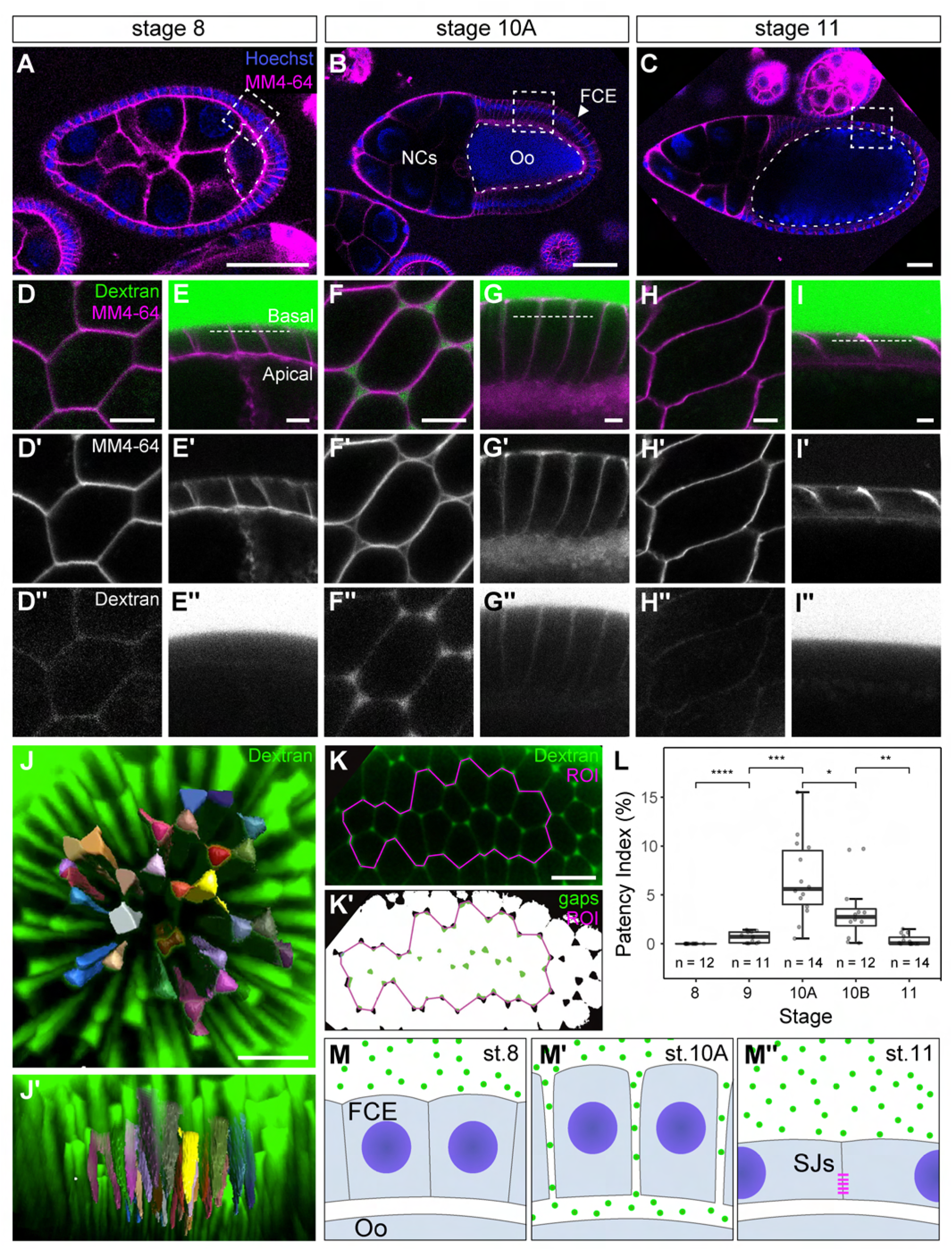
Transient opening of cell vertices enables paracellular transport through the follicle epithelium. (A-C) Confocal sections of living egg chambers at early (stage 8; A), mid- (stage 10A; B) and late (stage 11; C) vitellogenic stages. MM4-64 labels cell membranes (magenta), Hoechst 33342 labels nuclei (blue). Anterior is to the left. Nurse cells (NCs), oocyte (Oo) and Follicle cell epithelium (FCE) are indicated in (B), position of mainbody follicle cells is indicated by boxes in (A,B,C). (D-I) Living wild-type egg chambers were incubated in medium containing FITC-dextran (40 kDa; green) and MM4-64 membrane dye (magenta). Close-ups of mainbody FCs are shown in early (stage 8; D-E), mid- (stage 10A; F-G) and late vitellogenic (stage 11; H-I) egg chambers. (E, G, I) show cross-sections, (D, F, H) show sections taken perpendicular to the apical-basal axis at 25% of total epithelium height from the basal surface (indicated by dashed lines in E,G,I). Note that FCs at stage 8 are cuboidal with hexagonal outlines (D,E). Only little dextran signal is detectable at FC contacts (D”,E”). At stage 10A FCs are columnar in cross-section (G) and show a rounded outline in top view (F). Note dextran-filled intercellular spaces between FCs (F”,G”). At stage 11, FCs have flattened (H, I), intercellular spaces are closed (H’) and dextran is excluded from passing the epithelium (H”,I”). (J,J’) Top (J) and side (J’) views of confocal stack showing pyramidal shapes of dextran-filled channels spanning the width of the FCE at stage 10A. Surface-rendering of segmented channels is shown in color (see video S1). (K-L) For quantification of patency, sections acquired at 25% of total epithelium height from the basal surface (K) were converted into binary images by applying a threshold (K’). The fraction of open spaces (green areas) of the area of a region of interest (ROI, magenta) was plotted (patency index, PI; L). Analysis of PI in vitellogenic follicles (stages 8 to 11) revealed that patency starts at stage 9, peaks at stage 10A, and is terminated by stage 11. Sample size (n) indicates number of egg chambers analyzed (from at least 5 animals per stage). Wilcoxon test: * p ≤ 0.05, ** p ≤ 0.01, *** p ≤ 0.001, **** p ≤ 0.0001. (M-M”) Cartoons representing changes in the FCE during patency. FCE, follicle cell epithelium; Oo, Oocyte; SJs, septate junctions. See text for details. Scale bars: (A-C), 50 μm; (D-I), 5 μm; (J), 15 μm; (K), 10 μm.

In early vitellogenic follicles (stage 8; Fig. 1D-E) little dextran signal was detectable at the boundaries between the hexagonally shaped FCs (Fig. 1D”). By contrast, in mid-vitellogenesis (stage 10A), when FCs are columnar and their initially hexagonal outlines have rounded up (Fig. 1F-G), prominent dextran-filled intercellular spaces were visible at FC vertices (Fig 1F”). The dextran-filled channels were shaped like triangular pyramids, which were widest at the basal FCE surface and tapered towards the apical side (Fig. 1J; video S1). At stage 10A the channels spanned the entire width of FCE, with continuous dextran-filled lumina reaching from the basal side to the oocyte surface (Fig. 1G”,J), indicating that the channels were open *(“patent”)*, thus allowing paracellular transport across the FCE. We refer to this condition as *patency*, which was described in other insect species as a process in which the formation of channels between FCs allows transport of yolk proteins from the hemolymph across the FCE to the oocyte during vitellogenesis (Patchin and Davey, 1968; Pratt and Davey, 1972).

At stage 11, when FCs have flattened, the dextran-filled spaces disappeared and the FCE became impermeable for dextran (Fig. 1H-I). Analysis of the size of dextran-filled paracellular gaps as a fraction of the total FC area (patency index, PI; Fig. 1L) revealed that patency starts with small openings on the basal side at the onset of vitellogenesis in stage 9. The size of the gaps increased towards a peak at stage 10A, with a PI of 6% (n=14) and openings reaching a diameter of up to 1.8 μm on the basal side. After stage 10A the PI decreased to reach prepatency levels by stage 11. Thus, patency persists through stages 9 to 10B, corresponding to approximately 16 hours.

This time window coincides with the major phase of yolk uptake by the oocyte, suggesting that patency allows paracellular transport of yolk proteins from the hemolymph across the FCE to the oocyte surface. To test whether patency rendered the FCE permeable not only for dextran, but also for a physiological cargo, we performed analogous experiments using extracts from flies expressing yolk protein 3 tagged with monomeric red fluorescent protein (YP3-mRFP; 77 kDa). Like dextran, YP3-mRFP entered the inter-FC spaces and crossed the FCE in patent follicles (Suppl. Fig. 1), demonstrating that patency allowed paracellular transport of yolk protein across the FCE.

### Assembly of occluding junctions marks the termination of patency

To investigate how patency is terminated we examined the assembly of SJs using endogenously YFP-tagged SJ proteins. FCs did not exhibit discernible SJs until late vitellogenesis (Fig. 2A,B,D,E,G,H; Mahowald, 1972). However, concomitant with the end of patency, the SJ proteins Neuroglian-YFP (Nrg-YFP; Fig. 2A-C) and Lachesin-YFP (Lac-YFP; Fig. 2D-F) accumulated at apico-lateral membranes, and the TCJ proteins Gliotactin-YFP (Gli-YFP; Fig. 2G-I) and Anakonda (Aka; Suppl. Fig. 2A) accumulated at TCJs, suggesting that a tight paracellular barrier was established. Consistent with this notion, in stage 11 follicles dextran entered the basolateral space between FCs up to the border of the newly formed SJs, but was excluded from the more apical region occupied by the SJs (Fig. 2C,F,I). Together these findings show that barrier function of the FCE is developmentally regulated, with transiently increased paracellular permeability during patency at mid-vitellogenesis and the establishment of a tight SJ barrier after patency.

**Figure 2.**
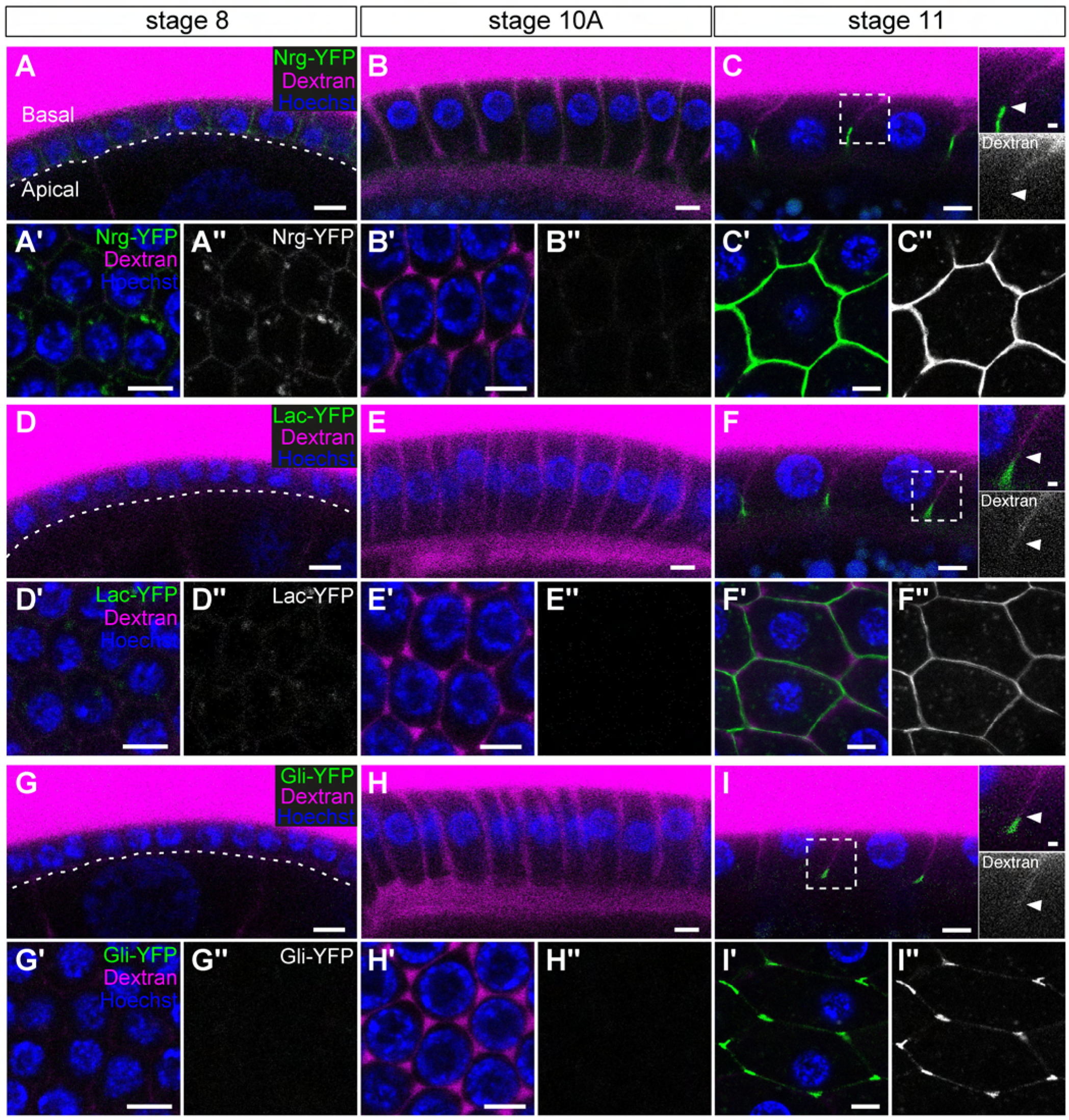
Assembly of occluding junctions marks the termination of patency. (A-I) Living egg chambers expressing YFP-tagged SJ proteins (green) Nrg-YFP^CPTI001714^ (AC), Lac-YFP^CPTI002601^ (D-F) or Gli-YFP^CPTI002805^ (G-I) were incubated in medium containing TRITC-dextran (10 kDa; magenta) and Hoechst 33342 (blue). Egg chambers are at early (stage 8; A,D,G), mid- (stage 10A; B,E,H) or late vitellogenic (stage 11; C,F,I) stage. Each panel shows a cross-section (top) and orthogonal view at the level of the nuclei (bottom) of mainbody FCs. Basal and apical surfaces of the follicle epithelium are indicated and the boundary between the follicle epithelium and the oocyte is marked by a dashed line in (A,D,G). Note that weak signals of Nrg-YFP and Lac-YFP are detectable at lateral membranes at stage 8 (A-A”; D-D”), disappear at stage 10A (B-B”; E-E”), and accumulate strongly at apico-lateral membranes at stage 11 (C-C”; F-F”). Gli-YFP is not detectable before stage 11 (G-I”) and accumulates at apico-lateral tricellular junctions at stage 11 (I-I”). Dextran is excluded from the FCE at stage 8 (A,D,G), enters intercellular spaces between FCs and the space between the FCE and the oocyte at stage 10A (B,E,H), and is again excluded from passage through the FCE at stage 11 (C,F,I). Close-ups of stage 11 FCs reveal that dextran penetrates intercellular spaces only basal to SJs marked by Nrg-YFP, Lac-YFP or Gli-YFP (C,F,I). Scale bars: (A-I), 5 μm; close-ups in (C,F,I), 1 μm.

To test whether TCJ assembly is required for termination of patency, we investigated mosaic egg chambers carrying clones of FCs lacking the TCJ protein Aka (Suppl. Fig. 2B). In *aka^L200^* loss-of-function clones Gli failed to accumulate at vertices and was instead distributed around the cell perimeter, indicating that TCJ formation in the FCE, as previously shown for other epithelia, depends on *aka* (Suppl. Fig. 2A-B; Byri et al., 2015). Interestingly, FC vertices within the *aka^L200^* clones appeared to be closed in post-patency follicles (Suppl. Fig. 2C). However, unlike wild-type controls, follicles bearing *aka* clones were permeable for 10 kDa dextran (Suppl. Fig. 2D). These findings suggest that TCJ assembly is not required for closing the intercellular gaps at the end of patency, but for sealing vertices to generate an impermeable FCE barrier.

### Sequential removal of adhesion proteins from vertices precedes the opening of intercellular channels

We focused on investigating the cellular processes that lead to the opening of FC vertices at stage 9. The opening and closing of the FCE barrier at cell vertices suggested that patency involves local remodeling of cell adhesion at TCJs. We therefore analyzed the distribution of cell-cell adhesion proteins prior to and during patency. Levels and distribution of the N-CAM orthologue Fas2, as detected using an endogenously tagged Fas2-YFP fusion protein or anti-Fas2 antibodies, changed dramatically before the onset of patency (Fig. 3; Suppl. Fig. 3; Gomez et al., 2012). In pre-vitellogenic egg chambers (stage 1-7) Fas2 was abundant at lateral FC membranes (Fig. 3A-B), while at stage 8, preceding the opening of intercellular spaces, Fas2 levels decreased markedly (Fig. 3C-D). Interestingly, however, Fas2 did not diminish uniformly around the cell perimeter, but receded selectively from vertices, while it remained detectable at bicellular contacts, resulting in a pattern of interrupted lines (Fig. 3C). Fas2 levels decreased further after intercellular channels opened, and became undetectable at maximum patency (stage 10A; Fig. 3E-F). When paracellular channels were sealed after patency (stage 11), Fas2 appeared again at FC-FC contacts, with elevated levels at vertices (Fig. 3G-H).

**Figure 3.**
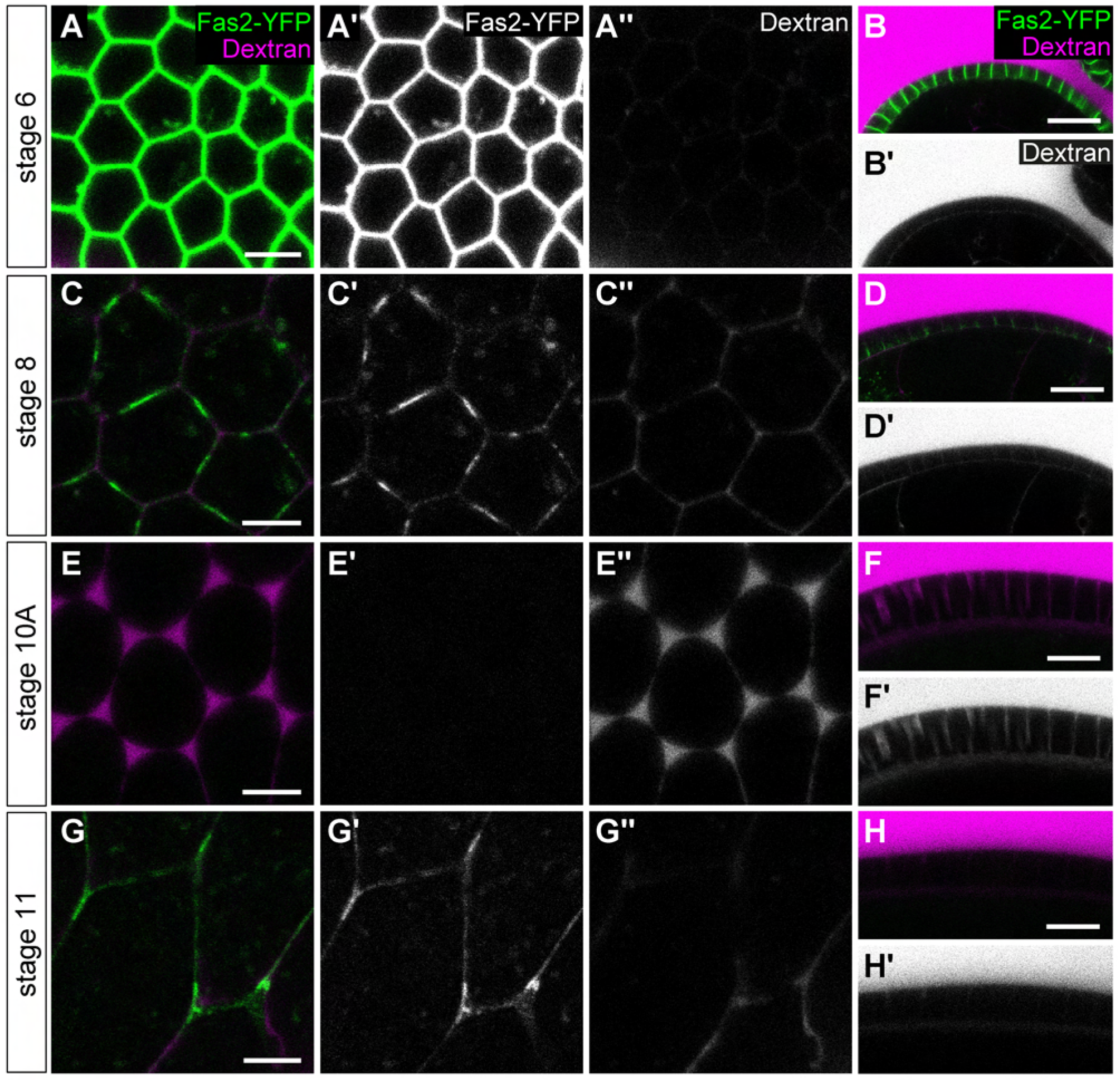
Removal of Fasciclin 2 from vertices precedes the opening of intercellular channels. (A-H) Living egg chambers from flies expressing *Fas2-*YFP^CPTI000483^, which labels all annotated Fas2 isoforms with YFP, were incubated in 10 kDa TRITC-dextran (magenta). (A,C,E,G) show top views, (B,D,F,H) show cross-sections. Fas2-YFP (green) localizes along cell perimeter at pre-vitellogenic stage 6 in top views (A,A’) and along the lateral membrane in cross-sections (B,B’). At stage 8, before the onset of patency, Fas2-YFP levels are strongly reduced overall and Fas2-YFP disappears from vertices (C,C’). Fas2-YFP is undetectable when intercellular channels are open during patency (stage 10A; E,F’). After patency (stage 11) Fas2-YFP re-appears and localizes to cell-cell contacts with increased levels at tricellular junctions (G,G’). Scale bars: (A,C,E,G), 5 μm; (B,D,F,H), 20 μm.

Similar to Fas2, *Drosophila* E-Cadherin (DE-Cad, referred to as E-Cad) and N-Cadherin (N-Cad) underwent dramatic changes just before patency (Suppl. Fig. 3). In pre-vitellogenic egg chambers, E-Cad and N-Cad accumulated at apical adhesion belts lining the perimeter of FCs and in addition were distributed along basolateral membranes (Suppl. Fig. 3A). Shortly before the onset of patency (stage 8), both E-Cad and N-Cad receded selectively from apical vertices, while they were maintained at bicellular contacts, resembling the distribution of Fas2 in the basolateral membrane (Suppl. Fig. 3F). At maximum patency (stage 10A) N-Cad completely vanished (Suppl. Fig. 3C; Tanentzapf et al., 2000), and, unlike Fas2, did not reappear after patency. By contrast, E-Cad remained detectable at bicellular junctions throughout patency, suggesting that E-Cad maintains bicellular adhesion between FCs during the process.

To further dissect the sequence of events that lead to patency we analyzed FC vertices in follicles expressing endogenously tagged versions of E-Cad (E-Cad-3xGFP or E-Cad-3x-mTagRFP) and the vertex-specific adhesion protein Sidekick (YFP-Sdk; Fig. 4). E-Cad was distributed in puncta along the basolateral membrane in addition to the apical AJs (video S2). At stage 6 both the basolateral E-Cad puncta and the apical AJs reached around the cell perimeter (Fig. 4A,B) and YFP-Sdk localized to tricellular vertices (Fig. 4C). At stage 8 E-Cad receded from vertices apically and slightly later basolaterally, while membranes at the vertex were still closed, indicating that removal of E-Cad precedes the opening of intercellular spaces (video S3). During stage 9 YFP-Sdk dispersed into multiple puncta along bicellular contacts, but was still detectable at vertices after E-Cad-GFP disappeared, suggesting that Sdk maintains adhesion at vertices in the absence of E-Cad. During patency (stage 10A) YFP-Sdk was absent and the separating cell membranes were bordered by E-Cad puncta in the basolateral membrane and by E-Cad bands at the apical side, suggesting that E-Cad-based adhesion limits the extent of paracellular openings. At stage 11 E-Cad and Sdk appeared again at the closed vertices. These findings suggest that the successive removal of E-Cad, N-Cad, Fas2 and Sdk from vertices is a precondition for patency.

**Figure 4.**
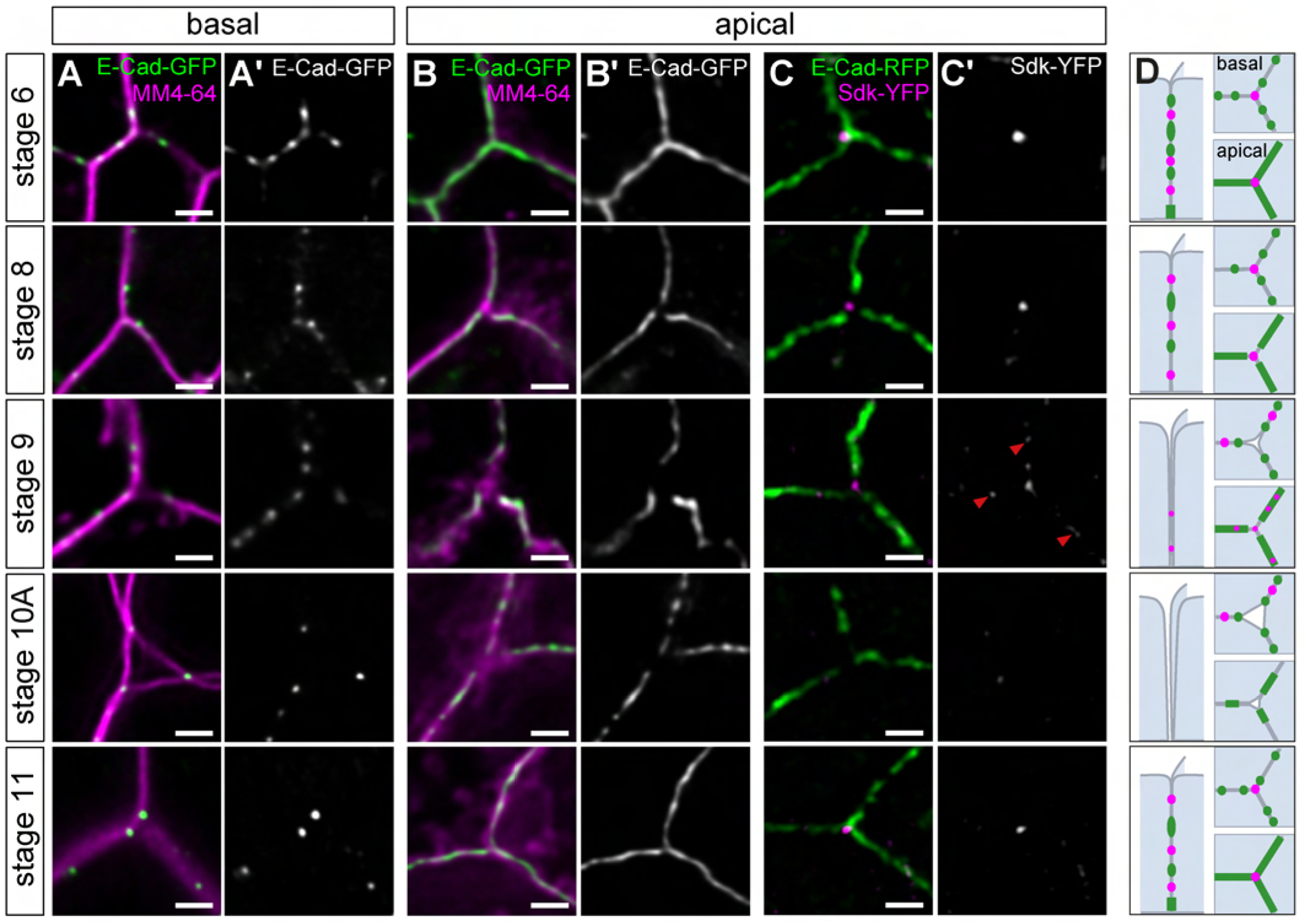
Sequential removal of E-Cadherin and Sidekick from vertices precedes patency. (A-B) Close-ups of single vertices in living egg chambers expressing E-Cad::3xGFP (green) labeled with MM4-64 membrane dye (magenta). (C) Close-ups of single vertices in living egg chambers expressing E-Cad::3x-mTagRFP (E-Cad-RFP; green) and YFP-Sdk^CPTI001692^ (magenta), which labels all annotated Sdk isoforms. FCs are shown in pre-vitellogenic (stage 6), early (stage 8), mid- (stages 9 and 10A) and late vitellogenic (stage 11) follicles. Sections of single vertices were acquired close to the basal (A) or close to the apical (B,C) surface of the FCE. Confocal images were processed using deconvolution. At pre-vitellogenic stage 6 ECad is distributed in puncta along the basolateral membrane (A) and in continuous adhesion belts near the apical surface (B), while YFP-Sdk accumulates at the vertex (C). E-Cad is removed from the vertex on the apical side by stage 8 and basally at stage 9, preceding the opening of intercellular channels. Note that at stage 8, when E-Cad has disappeared from the apical vertex, Sdk is still present. At the onset of patency (stage 9) Sdk signal decreases and Sdk puncta disperse into adjacent bicellular contacts. At stage 10A, single E-Cad puncta limit the separating membranes at the open intercellular spaces basally, while larger E-Cad patches limit the open vertices apically. Note that diffuse MM4-64 signal from the apical FCE and oocyte surface obscures a clear view of the intercellular space on the apical side. At stage 11, when intercellular spaces are closed, E-Cad puncta re-appear at the basolateral vertex and a continuous apical ZA, including the vertex, is restored apically. Sdk is absent from the vertex at stage 10A and re-appears after patency. (D) Cartoons representing the distribution of E-Cad (green) and Sdk (magenta) at basolateral and apical membranes, viewed from the side and from top, respectively, at the different stages. See text for details. Scale bars: 1 μm.

### Modulation of E-Cad-based adhesion at vertices regulates the extent of patency

Since E-Cad puncta limited the intercellular openings (Fig. 4M, video S3), we asked whether modulating E-Cad-based adhesion in FCs affects patency. Removing E-Cad function in *shg^1^* or *shg^2IH^* mutant clones was not informative, as the cells lacking E-Cad function were extruded from the FCE (not shown). We therefore used RNAi to partially deplete E-Cad from FCs. *GR1*-Gal4-driven expression of *E-Cad* dsRNA in FCs led to 8.4-fold (n=5, p=1.4×10^−6^) reduced ECad levels at FC junctions at stage 10A, but did not affect overall egg chamber morphology (Fig. 5B). Strikingly, however, at stage 10A the intercellular openings were enlarged compared to controls, and openings frequently merged at the basal side, resulting in the loss of adhesion not only at vertices, but also at bicellular contacts (Fig. 5B’). In mosaic follicles expressing *ECad* dsRNA in FC clones, this effect was limited to the *E-Cad* dsRNA expressing cells and did not spread beyond the boundary with adjacent wild-type cells (Fig. 5I), indicating a cell-autonomous role of E-Cad-based adhesion in patency. Surprisingly, despite the dramatic effect of E-Cad depletion on patency at stage 10A, the open intercellular spaces were closed again by stage 11 (Fig. 5B”), suggesting that residual levels of E-Cad, or the presence of other adhesion proteins, were sufficient for re-establishing cell contacts after patency.

**Figure 5.**
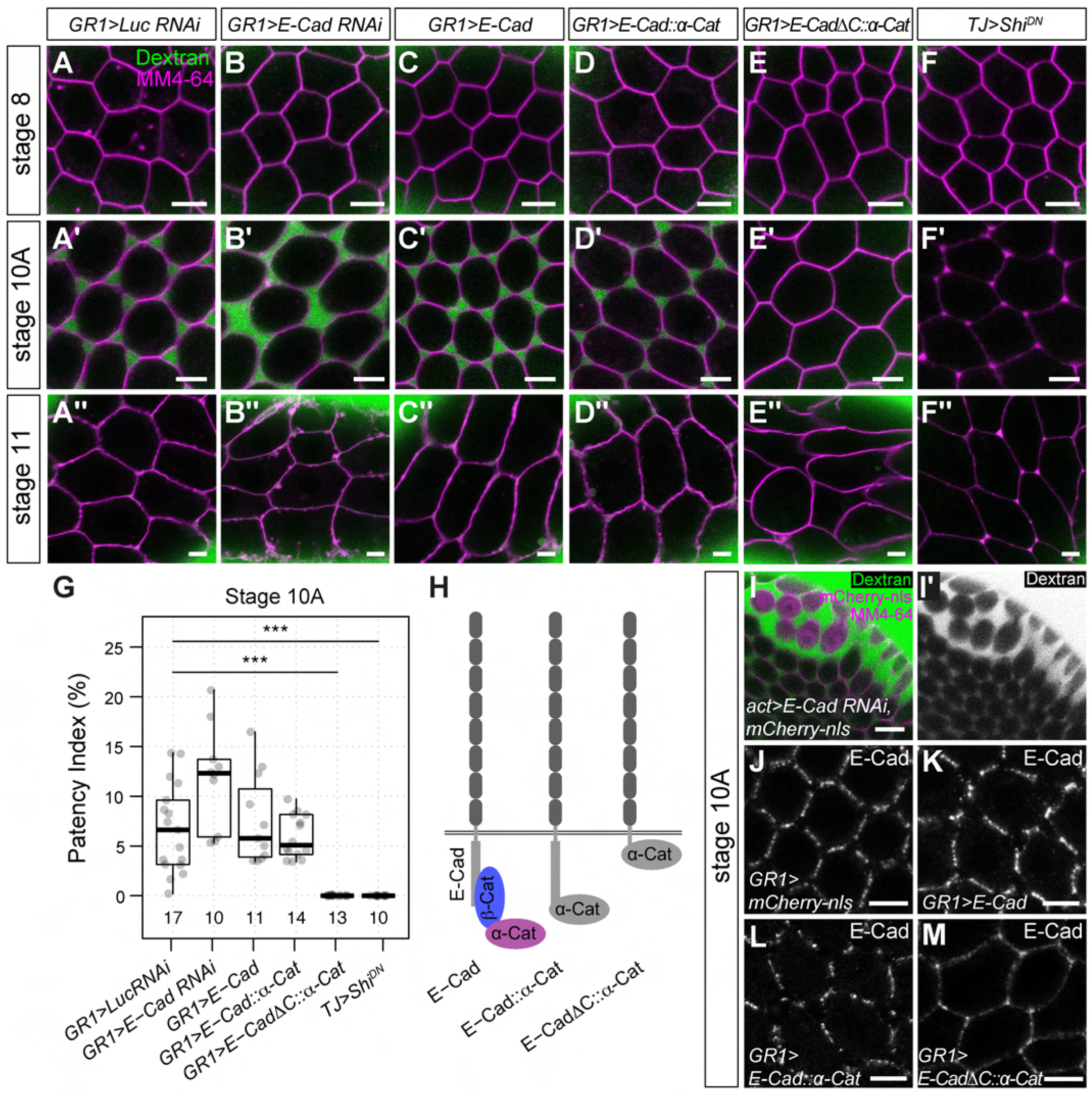
Modulation of E-Cad-based adhesion at vertices controls the extent of patency. (A-F) Living egg chambers were incubated with FITC-dextran (40 kDa; green) and MM4-64 membrane dye (magenta). FCs in early (stage 8), mid- (stage 10A) and late vitellogenic (stage 11) egg chambers are shown in sections acquired at 25% of total epithelium height. (A-E) *GR1-Gal4* was used to express *luciferase (luc)* dsRNA (A), *E-Cad* dsRNA (B), or to overexpress wild-type E-Cad (C), an E-Cadherin-α-Catenin fusion protein (E-Cad::α-Cat; D), or an E-Cad-α-Catenin fusion protein lacking the cytoplasmic domain of E-Cad (E-CadΔC::α-Cat; E). (F) *Traffic jam* (*tj*)-Gal4 was used to express dominant-negative dynamin (Shi-DN) in FCs after stage 6. Note that while opening and closing of vertices was unperturbed in the *luc* RNAi control (A-A”), RNAi-mediated depletion of E-Cad led to enlarged and partially merged openings at stage 10A (B’), which nonetheless closed at stage 11 (B”). Overexpression of wild-type E-Cad (C,C”) or of full-length E-Cad::α-Cat (D,D”) had no apparent effect on patency, whereas E-CadΔC::α-Cat expression completely blocked patency and led to abnormal cell shapes from stage 10A (E’,E”). Expression of dominant-negative dynamin (F) completely impeded the formation of dextran-filled channels at stage 10A (F’) and led to accumulations of MM4-64 signal at vertices from stage 10A (F’,F”). (G) Quantification of patency index at stage 10A. Sample size (number of egg chambers analyzed per genotype) is indicated at the bottom. Wilcoxon test: *** p ≤ 0.001. (H) Schemes of wild-type E-Cad, E-Cad::α-Cat and E-CadΔC::α-Cat fusion proteins. (I) E-Cad was depleted in a clone of FCs expressing *E-Cad* dsRNA and mCherry-nls. Note enlarged dextran-filled spaces (green) around *E-Cad* dsRNA-expressing cells, but not between adjacent wild-type cells. (J-M) Anti-ECad immunostaining reveals that wild-type E-Cad (J), overexpressed wild-type E-Cad (K) and E-Cad::α-Cat (L) are efficiently removed from vertices, whereas E-CadΔC::α-Cat escapes removal and accumulates at vertices (M). Scale bars (I), 10 μm; all other panels, 5 μm.

Aiming to increase E-Cad-based adhesion, we overexpressed E-Cad in FCs, resulting in 2.5-fold (n=6, p=1.3×10^−4^) higher E-Cad levels at FC-FC junctions (Fig. 5C,K). However, vertex-specific removal of E-Cad as well as patency appeared unperturbed in these egg chambers (Fig. 5C’,G,K). Overexpression of N-Cad gave corresponding results (data not shown). Therefore, to block removal of E-Cad from FC contacts prior to patency, we expressed in FCs a fusion protein consisting of DE-Cad lacking its cytoplasmic domain fused to α-Catenin (E-CadΔC::α-Cat), which was shown to stabilize AJs through constitutive linkage to the actin cytoskeleton (Pacquelet and Rørth, 2005). Whereas wild-type E-Cad was removed from vertices at stage 10A (Fig. 5K), E-CadΔC::α-Cat accumulated at vertices, indicating that the fusion protein was resistant to the vertex-specific removal mechanism (Fig. 5M). Strikingly, expression of E-CadΔC::α-Cat completely blocked the opening of vertices (Fig. 5E’,G), indicating that stabilizing AJs is sufficient to prevent patency. Interestingly, this effect was not observed upon expressing a full-length E-Cad-α-Catenin fusion protein containing the cytoplasmic domain of DE-Cad (E-Cad::α-Cat; Fig. 5D). Unlike E-CadΔC::α-Cat, full-length E-Cad::α-Cat was removed from vertices and did not affect patency (Fig. 5D’,G,L), indicating that the cytoplasmic domain of E-Cad is required for removal from vertices, possibly through endocytic turnover (Woichansky et al., 2016). Taken together, these findings show that vertex-specific removal of E-Cad-based adhesion is necessary for initiating patency and that changes in the levels of E-Cad modulate the extent of intercellular openings.

### Endocytosis is required for opening of vertices

To further probe the mechanism underlying the initiation of patency we tested the requirement of endocytosis by expressing in FCs a dominant-negative form of the GTPase dynamin (Shi-DN; Moline et al., 1999), which blocks the scission of endocytic vesicles from the plasma membrane. Indeed, expression of Shi-DN completely blocked the opening of FC vertices and led to conspicuous vertex-specific accumulations of MM4-64 signal (Fig. 5F, G), suggesting a special requirement for endocytosis to maintain membrane homeostasis at FC vertices. These findings are consistent with the idea that patency depends on endocytic turnover of adhesion proteins, including E-Cad and Fas2.

### Removal of Fas2 is required for patency

We next tested whether the removal of Fas2 (Fig. 3) is required for patency. We overexpressed in the FCE two different Fas2 transmembrane isoforms either containing or lacking a cytoplasmic PEST motif with a potential role in protein degradation (Lin et al., 1994). Immunostainings revealed that *GR1*-Gal4-driven overexpression of Fas2(PEST-) or Fas2(PEST+) isoforms led to strong accumulation of Fas2 protein at basolateral FC membranes at stage 10A (Fig. 6F; not shown for Fas2(PEST+), when Fas2 was absent in wild-type follicles (Fig. 6E; Gomez et al., 2012). This suggests that the mechanism of Fas2 degradation in FCs can be overcome by Fas2 overexpression, and that the PEST motif is not required for Fas2 degradation in this situation. To assess the effect of Fas2 overexpression on patency we analyzed the permeability of egg chambers for dextran (Figure 6A-C). Strikingly, overexpression of either Fas2(PEST+) or Fas2(PEST-) completely blocked the opening of vertices and the rounding-up of FCs compared to controls at stage 10A (Figure 6B’, C’, D), indicating that Fas2 needs to be absent to allow patency.

**Figure 6.**
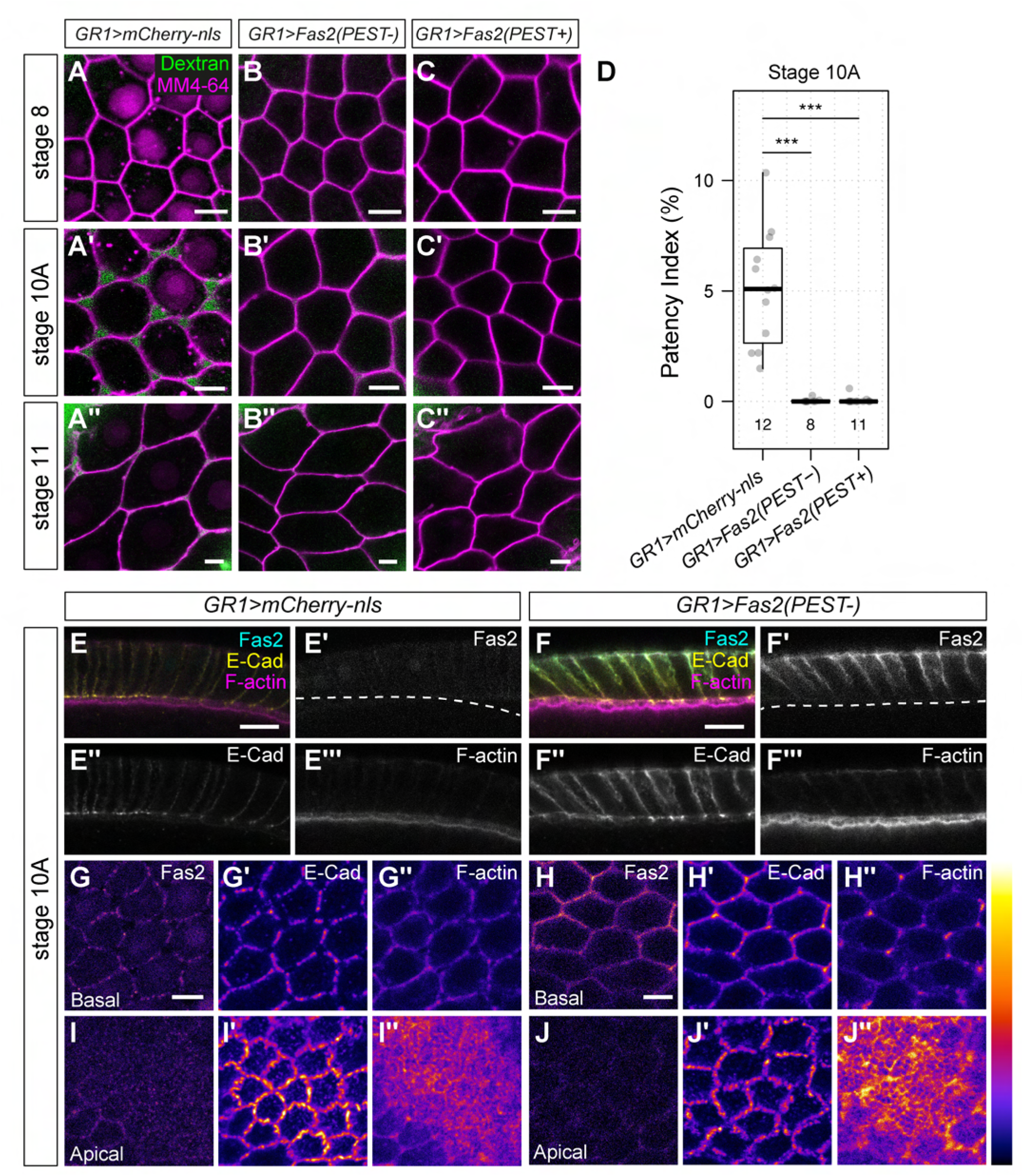
Fasciclin 2 controls actin organization and adhesion at vertices. (A-C) Living egg chambers from flies overexpressing Fas2(PEST-) (B) or Fas2(PEST+) (C) isoforms, or expressing mCherry-nls as control (A) incubated with FITC-dextran (40 kDa; green) and MM4-64 membrane dye (magenta). At stage 8, Fas2(PEST-) (B) or Fas2(PEST+) (C) overexpression has no apparent effect on FC morphology compared to the control (A). At stage 10A, control FCs have rounded up and show dextran-filled intercellular spaces (A’), whereas Fas2(PEST-) (B’) or Fas2(PEST+) (C’) overexpressing FCs retain their hexagonal shapes and fail to form intercellular spaces (B’,C’). (D) Quantification of patency index at stage 10A. The number of egg chambers analyzed per genotype is indicated at the bottom. Wilcoxon test: ***p ≤ 0.001. (E-J) Anti-Fas2 immunostaining of control (mCherry-nls; E,G,I) and Fas2(PEST-) overexpressing (F,H,J) stage 10A follicles. Cross-sections (E,F) show that while Fas2 is not detectable in control FCs at stage 10A (E’), Fas2 overexpression leads to strong accumulation of Fas2 (F’) and elevated E-Cad levels in the basolateral membrane, colocalizing with Fas2 (E”,F”). (G-J) En-face views of sections taken near the basal (G,H) or apical (I,J) surface. Note that E-Cad and F-actin are removed from vertices in the control (G’,G”), but accumulate on basolateral vertices upon Fas2(PEST-) overexpression (H’,H”). Overexpression of Fas2(PEST+) gave corresponding results (not shown). Fluorescence intensities are color-coded using the heat map shown to the right. Scale bars: (A-C) and (G-J), 5 μm; (E-F), 10 μm.

### Fas2 overexpression is sufficient to maintain E-Cad and F-actin at vertices

Surprisingly, we found that in Fas2(PEST-) or Fas2(PEST+) overexpressing egg chambers ECad and F-actin signals at basolateral FC membranes were increased compared to controls (Fig. 6F; data not shown for Fas2(PEST+). This was particularly evident at vertices, where ECad and F-actin accumulated when Fas2 was overexpressed, whereas vertices were depleted from E-Cad and F-actin in controls (Fig. 6G-H”). By contrast, on the apical side, where overexpressed Fas2 did not accumulate (Fig. 6F’), E-Cad was still removed from vertices (Fig. 6J’), suggesting that distinct mechanisms control E-Cad removal from the basolateral and apical parts of the vertex, respectively. Together these findings suggest that Fas2, in addition to mediating cell adhesion by itself, may act to promote or maintain E-Cad accumulation at basolateral FC vertices, possibly through organizing the actin cytoskeleton.

### Cortical actin is remodeled prior to patency initiation

To determine whether the removal of E-Cad from vertices is correlated with changes in the actomyosin cytoskeleton, we analyzed the distribution of E-Cad, F-actin and the GTPase Rho1, which induces actomyosin contractility through activating Rho-kinase and inhibiting myosin phosphatase, at the onset of patency (Figure 7A-F). Preceding patency (stage 7) Factin and E-Cad accumulated at basolateral vertices, while Rho1 was enriched at lateral membranes around the cell perimeter (Fig. 7A-C). At the onset of patency (stage 9), the accumulation of Rho1 at the membrane decreased (Fig. 7D). At the same time cortical F-actin was reduced (Fig. 7B,E), E-Cad disappeared from basal vertices (Fig. 7C,F), and the apical ECad-positive AJs became corrugated, suggesting that cortical tension decreased (Fig. 7C’,F’).

**Figure 7.**
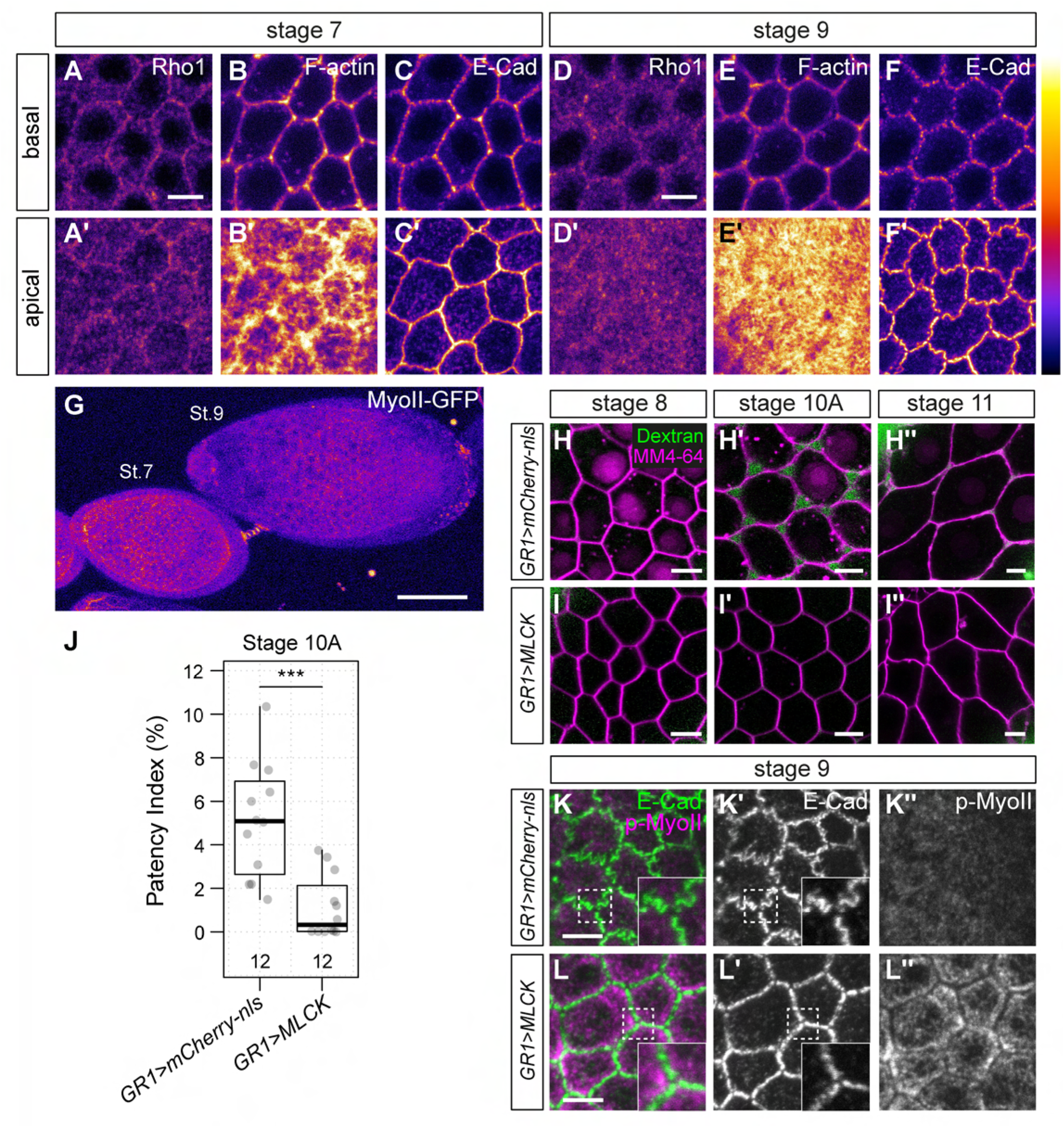
Increased actomyosin contractility prevents patency. (A-F) Immunostaining of Rho1 (A,D), F-actin (phalloidin; B,E) and E-Cad (C,F) in wild-type egg chambers. Sections taken close to the basal (A-F) and apical (A’-F’) side are shown. Fluorescence intensities are color-coded using the heat map shown to the right. In pre-vitellogenic (stage 7) follicles Rho1 and F-actin are enriched at the cortex along the cell perimeter (A,B), and F-actin (B) and E-Cad (C) accumulate at basolateral vertices. At stage 9, Rho1 cortical enrichment is lost (D), basolateral F-actin signals decrease (E) and E-Cad is removed from vertices (F,F’). Note that E-Cad-marked adherens junctions are corrugated at the apical side, consistent with a decrease in cortical tension (compare C’ and F’). (G) Living egg chambers expressing Myosin II-GFP (MyoII-GFP). Note the decrease in MyoII-GFP levels from stage 7 to stage 9. (H-I) Living egg chambers expressing mCherry-nls (control; H) or activated myosin light chain kinase (MLCK; I) were incubated in medium containing FITC-dextran (40 kDa; green) and MM4-64 membrane dye (magenta). Note that MLCK overexpression blocks the formation of dextran-filled intercellular spaces at stage 10A (I,I’). (J) Quantification of patency index at stage 10A. Number of egg chambers analyzed per genotype is indicated at the bottom. The control *(GR1>mCherry-nls)* data is the same as in *UAS-Fas2* overexpression experiments (Fig. 6D), which were done in parallel. Wilcoxon test: ***p ≤ 0.001. (K-L) Immunostaining of E-Cad and phospho-Myosin II (p-MyoII) in mCherry-nls control (K) and MLCK-overexpressing (L) egg chambers. Phospho-Myosin II signals are elevated in MLCK overexpressing FCs (L”) compared to control (K”). Note that while adherens junctions are corrugated in the control (K’), they are straight in MLCK-overexpressing FCs (L’). Scale bars: (G), 50 μm; all other panels, 5 μm.

### Actomyosin contractility regulates opening of vertices

To corroborate these findings, we investigated the levels of myosin II-GFP (myoII-GFP) in living egg chambers throughout oogenesis. MyoII-GFP signals in FCs decreased from stage 7 to stage 9 (Fig. 7G), concomitant with decreasing accumulation of Rho1 at FC membranes (Fig. 7A,D), suggesting that cortical tension in FCs is decreasing during this interval. This notion was consistent with decreasing levels of the tension-sensitive protein vinculin (vinculin-GFP; Maartens et al., 2016) at FC junctions from stage 6 to stage 9 (data not shown) and with direct measurements of cortical tension by laser ablation experiments (Balaji et al., 2019). We hypothesized that a decrease in cortical tension could play a role in initiating patency. To test this idea we over-activated myosin II by expressing a constitutively activated form of chicken myosin light chain kinase (MLCK; Kim et al., 2002) in FCs under the control of *GR1-Gal4* (Fig. 7I). Indeed, MLCK expression led to a prominent increase in myoII activation, as detected by anti-phospho-Myosin II staining, at the apical side of FCs in stage 9 egg chambers compared to controls (Fig. 7K-L). This was associated with a conspicuous change in the morphology of AJs, which were straight in control FCs, but corrugated in MLCK-expressing FCs (Fig 7K’,L’), suggesting that MLCK-induced myosin contractility led to an increase in cortical tension, consistent with the findings of (Balaji et al., 2019). Strikingly, FCs in MLCK-expressing follicles failed to round up and instead maintained their hexagonal shape at stage 10A, and showed a strong reduction in the size of intercellular openings (Fig. 7H-J). Expression of the catalytic domain of Rho-kinase (Rok^CAT^; Winter et al., 2001) as an independent means to increase actomyosin contractility led to comparable effects (data not shown). These findings suggest that a decrease in cortical tension prior to the onset of patency is necessary for the subsequent cell rounding and opening of intercellular spaces. Thus, removal of adhesion and reduction of cortical tension appear to act in concert to promote the opening of vertices during patency.

## Discussion

We described the cellular events and mechanisms underlying patency, a developmentally regulated process in which the dynamic remodeling of cell vertices leads to dramatic changes in paracellular permeability of an epithelial barrier. Patency offers a powerful new paradigm, accessible to live imaging, genetic analysis, and pharmacological manipulation, for studying the mechanisms of epithelial barrier remodeling *in vivo*.

First, we show that during patency the paracellular barrier of the follicle epithelium is breached at tricellular vertices, opening up intercellular channels that allow transport of proteins across the epithelium for uptake by the oocyte. Second, patency is a transient process initiated by the sequential removal of multiple adhesion proteins from tricellular vertices, preceding their polarized basal-to-apical opening, and terminated by the assembly of tricellular occluding junctions that restore a tight paracellular barrier. Third, we show that modulation of E-Cad- or Fas2-based adhesion and of actomyosin contractility regulates the extent of opening of intercellular spaces, and that patency can be prevented by artificially stabilizing adherens junctions. Thus, we defined the cellular events and underlying mechanisms that lead to the transient opening of the FCE paracellular barrier, a crucial step for yolk uptake during oogenesis.

Our findings reveal a key role of cell vertices as gateways controlling paracellular transport across epithelia, and demonstrate that the dynamic interaction between adhesion molecules and the cytoskeleton at vertices governs epithelial permeability. We showed that lateral (Fas2, E-Cad), apical/AJ-associated (E-Cad, N-Cad) and vertex-specific (Sdk) adhesion proteins are removed from FC vertices in a characteristic sequence of events prior to the onset of patency. The spatiotemporal pattern of removal of these proteins correlates with the polarized basal-to-apical opening of paracellular channels, suggesting that the successive loss of adhesion plays a permissive role in the process. This implies that the relevant adhesion molecules are regulated differentially along the apical-basal axis. In particular, we showed that the differential behavior of apical *(zonula adherens)* and basolateral pools of E-Cad, as well as the levels of E-Cad protein, determine the spatiotemporal pattern and the size of intercellular openings. The NCAM orthologue Fas2, by contrast, appears to act at a higher regulatory level, as overexpression of Fas2 transmembrane isoforms led to accumulation of E-Cad and F-actin at basolateral vertices, suggesting that besides providing adhesion, Fas2 modulates the actin cytoskeleton as well as turnover of E-Cad in FCs. This could be mediated by a function of the Fas2 cytosolic domain in anchoring F-actin, possibly via Ankyrin (Pielage et al., 2008). Alternatively, Fas2 overexpression may have indirect effects, *e.g*. on the endocytosis machinery. Prolonged Fas2 expression led to a phenotype resembling previtellogenic stages, suggesting that Fas2 removal represents a key step in the sequential loss of adhesion leading to patency. Fas2 endocytosis is induced by the Ser/Thr kinase Tao and reduces lateral adhesion between FCs, leading to shrinking of lateral membranes (Gomez et al., 2012). Interestingly, the onset of Fas2 removal at stage 6 coincides with the mitotic-to-endocycle switch in FCs orchestrated by Notch signaling (Deng et al., 2001; Lopez-Schier and St Johnston, 2001). Among the cell cycle regulators downstream of Notch is the anaphase-promoting complex/cyclosome (APC/C) co-activator Fizzy-related (Fzr), which allows G1 progression and promotes endocycles (Shcherbata et al., 2004). Intriguingly, Fzr was reported to be required for removal of Fas2 in glia cells (Silies and Klambt, 2010). This suggests the intriguing possibility that the Notch-induced cell cycle switch in FCs is linked to Fas2 degradation, which in turn may reduce anchorage of F-actin and consequently promote E-Cad turnover, eventually leading to the opening of FC vertices.A fascinating feature of patency is the locally restricted removal of adhesion molecules from cell vertices. The underlying mechanism is not yet clear. In principle, endocytic uptake, recycling, degradation, or lateral mobility of adhesion molecules could be differentially regulated between bi- and tricellular contacts, possibly mediated by a specialized membrane trafficking machinery at vertices. While plant cells contain a cell-corner-specific membrane compartment marked by the GTPase Rab-A5c (Kirchhelle et al., 2016), we found no evidence for vertex-specific vesicles by examining the subcellular distribution of 26 different YFP-labeled Rab GTPases in FCs (data not shown). Our results support the idea that E-Cad turnover at vertices is promoted by specific properties of the actomyosin network at these sites. Consistent with this notion, actin filaments in vertebrate cells are anchored in an end-on fashion at tricellular AJs, as opposed to a side-on association at bicellular AJs, resulting in a distinct distribution of mechanical tension at vertices (Yonemura, 2011). Moreover, Myosin II-dependent tensile and shear stress, respectively, were found to have opposite effects on the levels of E-Cad during junctional remodeling (Kale et al., 2018), and inhibiting Myosin II activity was shown to reduce E-Cad enrichment and dynamics at vertices in the *Drosophila* embryonic ectoderm (Vanderleest et al., 2018). Tension at vertices could also modulate endocytic turnover of E-Cad through tyrosine phosphorylation by Src kinases, which can be activated by mechanical force (Wang et al., 2005). Thus, the geometry of tricellular vertices, and the resulting distribution of tension at these sites (Higashi and Miller, 2017), may account for their distinct properties with respect to junctional remodeling.

Interestingly, we found that concomitant with the loss of adhesion at lateral membranes between stages 7 and 9, cortical actin, Rho1 and MyoII levels decrease. Rho1-GTP activates the kinase Rok, which in turn phosphorylates Myosin-II regulatory light chain (MRLC) and thereby induces actomyosin contractility in many morphogenetic processes (Lecuit and Lenne, 2007). However, our findings indicate that during vitellogenesis cortical actomyosin contractility in FCs is decreasing rather than increasing, despite forces arising from the growth of the underlying oocyte. This is consistent with recent work demonstrating that junctional tension in FCs decreases during oogenesis, explaining the appearance of corrugated FC junctions at stage 9 (Balaji et al., 2019). Accordingly, we show that increased actomyosin contractility upon MLCK overexpression impairs cell rounding and blocks the opening of intercellular spaces. The finding that cortical tension decreases as FCs round up was unexpected, since cell rounding in isolated cells, *e.g*. at the onset of mitosis, was shown to be associated with an increase in cortical tension (Stewart et al., 2011). However, cortical tension also modulates the strength of adhesion by regulating the recruitment, recycling, and lateral mobility of adhesion complexes via mechanical feedback (Hoffman and Yap, 2015). Accordingly, VE-Cadherin recruitment and AJ size were shown to increase or decrease with myosin II activation or inhibition, respectively, in endothelial cells (Liu et al., 2010). Generally, in the absence of adhesion to neighboring cells or to a substrate, cells assume a spherical shape. Hence, we propose that reduction of cortical tension in FCs weakens AJs and permits the redistribution of basal E-Cad puncta to bicellular junctions, thus reducing adhesion at vertices and allowing cells adopt rounded shapes. Reduced actomyosin contractility at the onset of patency may also initiate the redistribution of Sdk from vertices to bicellular junctions, as Sdk interacts directly with MRLC (Uechi and Kuranaga, 2019) and accumulation of Sdk at vertices depends on active myosin II (Letizia et al., 2019). Thus, decreasing cortical tension and sequential loss of adhesion appear to act in concert to promote the opening of intercellular spaces between FCs.

In different insects orders patency was reported to be regulated by Juvenile Hormone (JH; Santos et al., 2019). In *Locusta migratoria* ovaries, JH triggers a G protein-coupled receptor signaling cascade leading to Na^+^/K^+^-ATPase activation, which in turn changes the ionic balance of FCs and causes cell shrinkage, promoting the formation of intercellular channels (Jing et al., 2018). This implies that, in addition to reduction of actomyosin contractility and the loss of cell adhesion at vertices, hormonally controlled modulation of cell volume might be involved in patency. While we did not find evidence supporting a role of JH signaling in initiating patency, a recent study reported a requirement for signaling by the steroid hormone Ecdysone in patency in *Drosophila* (Row and Deng, 2020). Intriguingly, in the mammalian testis, testosterone promotes the opening of the blood-testis barrier through modification of TJs between Sertoli cells (Chakraborty et al., 2014). It will be fascinating to study how hormones might regulate FC junction remodeling during patency.

Follicle patency is a conserved feature of oogenesis in insects (Raikhel and Dhadialla, 1992). Interestingly, vitellogenesis in oviparous vertebrates (fish, amphibians, reptiles, birds, monotremes) resembles the process in insects: the yolk protein precursor vitellogenin is synthesized in the liver, secreted into the blood, and taken up into the oocyte via receptor-mediated endocytosis (Romano et al., 2004). Intriguingly, patent intercellular channels between follicle cells, and passage of tracers or of bloodborne proteins through these channels, were described in eggs of fish (Wallace and Selman, 1990), amphibians (Dumont, 1978) and reptiles (Neaves, 1972), suggesting that patency is conserved in oviparous animals.

A growing body of work highlights the key roles of tricellular vertices in epithelial barrier remodeling processes in health and disease (Higashi and Miller, 2017; Bosveld et al., 2018). Endothelial cell vertices are preferential routes for leukocyte transmigration in blood vessels (Burns et al., 2003), and bacterial pathogens, such as Group A *Streptococcus* (Sumitomo et al., 2016), use TCJs as entry paths to cross epithelial barriers during infection. As these processes involve the dynamic remodeling of endothelial or epithelial cell vertices, it will be exciting to explore whether the underlying mechanisms that mediate the opening and closing of vertices are similar to those that mediate follicle cell patency in *Drosophila*.

## Materials and Methods

### *Drosophila* strains and genetics

The following *Drosophila* stocks are described in FlyBase and were obtained from the Bloomington stock center, unless noted otherwise: *Act5C-Gal4* (4414), *GR1-Gal4* (36287), fat body (FB)-Gal4 (Werthebach et al., 2019), traffic jam-Gal4 (tj-Gal4; gift from Veit Riechmann, University of Heidelberg, Germany), *Act5C>CD2>Gal4 UAS-mRFP-nls* (30558), UAS-mCherry-nls, UAS-*luc* RNAi (31603), *UAS-E-Cad* RNAi (VDRC-27081), *UAS-E-Cad* RNAi (NIG-3722R-1), UAS-DE-Cad (58494; Pacquelet and Rørth, 2005), UAS-*E-Cad::αCat* (65580), *UAS-E-CadΔC::αCat* (67415), UAS-Fas2A(PEST+), UAS-Fas2A(PEST-) (Silies and Klambt, 2010), *Gli*-YFP^CPTI002805^, *La*c-YFP^CPTI002601^, *Nrg*-YFP^CPTI001714^, *YFP-Sdk^CPTI001692^, Fas2-* YFP^CPTI000483^ (Lye et al., 2014), E-Cad::3xGFP, E-Cad::3xmTagRFP, MyoII::3xGFP (Pinheiro et al., 2017), Vinculin-GFP (Maartens et al., 2016), UAS-Shi-DN (5811), UAS-MLCK (37528; Kim et al., 2002), UAS-Rok.CAT (6669; Winter et al., 2001), *hs-Flp^122^, aka^L200^ FRT40A* (Byri et al., 2015), *FRTG13 shg^1^*, *FRTG13 shg^2IH^, ubi-nlsGFP FRT40A* (5189), *FRTG13 ubi-nlsGFP* (5826). Oregon R was used as the wild-type strain.

### Generation of marked clones in the follicle epithelium

To generate mosaic animals carrying clones of *aka* homozygous mutant cells we crossed *y w hs-Flp^122^; ubi-nlsGFP FRT40A* females to *aka^L200^ FRT40A* males. Mosaics carrying *DE-Cad (shg)* mutant clones were generated correspondingly using *FRTG13 shg^1^* or *FRTG13 shg^2IH^* and *FRTG13 ubi-nlsGFP* chromosomes. To generate flip-out clones expressing *UAS-E-Cad* RNAi, we crossed *y w hs-Flp^122^; UAS-E-Cad RNAi* females to *Act5C>CD2>Gal4, UAS-mRFP-nls* males. Pupae (48-72 hours after puparium formation) were heat-shocked for 30 min at 37°C to induce Flp-mediated recombination.

### Molecular biology

UAS-YP3-mRFP was generated as follows: The yolk protein 3 (CG11129) coding sequence (420 aa; GenBank AAF48314.2), fused C-terminally to a Streptavidin-binding peptide sequence, a (GGS)4 linker sequence, and the mRFP coding sequence, was synthesized (GenScript), cloned into pUASt-attB (EcoRI/XbaI), and integrated into the *attP40* and *attP2* landing sites using PhiC31 integrase (Bischof et al., 2007).

### Antibodies and Immunostainings

Adult females collected 24-48h after eclosion were fed on yeast at 25°C for two days before dissecting ovaries in M3 insect medium (Sigma). Egg chambers were transferred into 4% paraformaldehyde in M3 medium/PBS and fixed at room temperature for 20 min. Primary antibodies were rat anti-DCAD2 (1:100; DSHB), rat anti-N-Cad DN-Ex #8 (1:100; DSHB), rabbit anti-Aka (1:500; Byri et al., 2015), mouse anti-Gli 1F6 (1:200; Schulte et al., 2003), mouse anti-Fas2 1D4 (directed against the Fas2 cytoplasmic domain; 1:20; DSHB) and rabbit anti-phospho-Myosin light chain 2 (1:50; Cell Signaling Technology). Goat secondary antibodies were conjugated with Alexa Fluor 488, 568 or 647 (1:500; Life Technologies). Factin was stained with TIRTC-or FITC-Phalloidin (1:1000; Sigma) added together with secondary antibodies. Egg chambers from wild-type flies expressing mCherry-nls were used as internal controls processed in the same reaction alongside with experimental genotypes.

### Microscopy and image analysis

Samples were imaged on a Leica SP8 confocal microscope with 40x/1.3 NA and 60x/1.4 NA oil immersion objectives. Where indicated, deconvolution was performed using the Leica Application Suite X software 3.5.1 (adaptive mode). Images were processed with FIJI (v2.0.0; Schindelin et al., 2012). OMERO.web 5.5.1 was used to adjust brightness and contrast. OMERO.figure 4.0.2 was used for assembling image panels. Figures were created in Adobe Illustrator CS6 (v.16.0.0). Imaris (v.9.0.0, Bitplane) was used to create 3D animations.

### Quantification of immunofluorescence signals

Efficiencies of E-Cad knockdown or overexpression were assessed based on anti-E-Cad immunostainings. Single confocal sections were acquired at the apical side of at least five egg chambers per genotype. Intensity of junctional E-Cad signals and of intracellular (background) signals was measured using ROIs of equal sizes. For each egg chamber, the mean of background was subtracted from the mean of junctional signal. Significance was determined using Student’s t-test.

### Dextran permeability assay and culture of egg chambers

Adult females collected 24-48h after eclosion were fed on yeast at 25°C for two days before dissecting ovaries in Shields and Sang M3 insect medium (Sigma) containing 10% FBS (HyClone; GE Healthcare) and 1x penicillin/streptomycin (from 100x stock; Thermo Fisher). Egg chambers were transferred into 200 μl of this medium supplemented with FITC-, TRITC-, or Alexa-647-labeled dextran (40 or 10 kDa respectively; 12.5 μg/ml; Sigma). Where indicated, membrane dye MM4-64 (2 nM; Santa Cruz Biotechnology) and Hoechst 33342 (1 μg/ml; Sigma) were added to label cell membrane and nuclei, respectively. For measuring FCE permeability egg chambers were mounted on glass slides with coverslips and imaged immediately after preparation.

For longer-term culture, follicles were kept in 8-well glass-bottom chambers (VWR 734-2061) with 200 μl M3 medium per well. To immobilize follicles on the glass surface, the glass-bottom chambers were coated with poly-D-Lysine (0.1 mg/ml in PBS; Sigma) for 1 h at 37°C before use. To induce osmotic shocks, 50 μl of 2.5x PBS (10x PBS diluted in M3 medium) were added during imaging to egg chambers cultured in 200 μl M3 medium to yield a final concentration of 0.5x PBS. Confocal stacks were acquired for one hour at 5-second intervals.

### Fly extract experiments

Fly extract was prepared according to https://dgrc.bio.indiana.edu/cells/modencode/Protocol-CME-W1-C18. Female flies expressing UAS-YP3-mRFP under the control of *GR1*-Gal4 or Oregon R flies (control) were frozen at −20°C and homogenized in M3 insect medium (300 flies in 6.8 ml). The homogenate was centrifuged at 1.500 x g at 4°C for 15 minutes. The pellet was discarded and the supernatant was incubated at 60°C for 5 minutes to inactivate tyrosinase. The sample was centrifuged again at 1.500 x g at 4°C for 90 minutes and the supernatant was collected. Egg chambers were incubated in this fly extract for imaging.

### Quantification of patency index and statistics

Confocal sections of mainbody FCs in the middle of the egg chamber (up to stage 8) or overlying the oocyte (after stage 8) were acquired at 25% of total epithelium height from the basal surface. A ROI including 10-15 cells (stages 8 and 10A) or 5-8 cells (stage 11) was selected in FIJI. After applying a mean filter the images were converted to binary mode using a manually selected threshold to represent the open intercellular spaces without bicellular junction signals. Within a given experiment the threshold was set in the control and applied to the experimental samples. The total area of all intercellular spaces (gaps) within the ROI was subtracted from the total area of the ROI to yield the cellular area. The patency index (PI) was calculated as the area of gaps divided by the cellular area (expressed in percent). Calculations, statistics and plots were performed in R (v.3.5.1). Since the data did not follow a normal distribution, significance was determined using the pairwise two-tailed Wilcoxon-Mann-Whitney-U test. P-values were adjusted for multiple testing using Holm’s method (Holm, 1979).

## Acknowledgements

We thank Yohanns Bellaiche, Mathias Beller, Christian Klämbt, Veit Riechmann, and Pernille Rorth for providing fly stocks and reagents. We thank Wilko Backer for expert technical help, Mylène Lancino for movie S1, Raphael Schleutker for help with data analysis and statistics, and Timo Betz and Veit Riechmann for discussions and comments on the manuscript.

## Supplementary Figure Legends

**Supplementary Figure 1.**
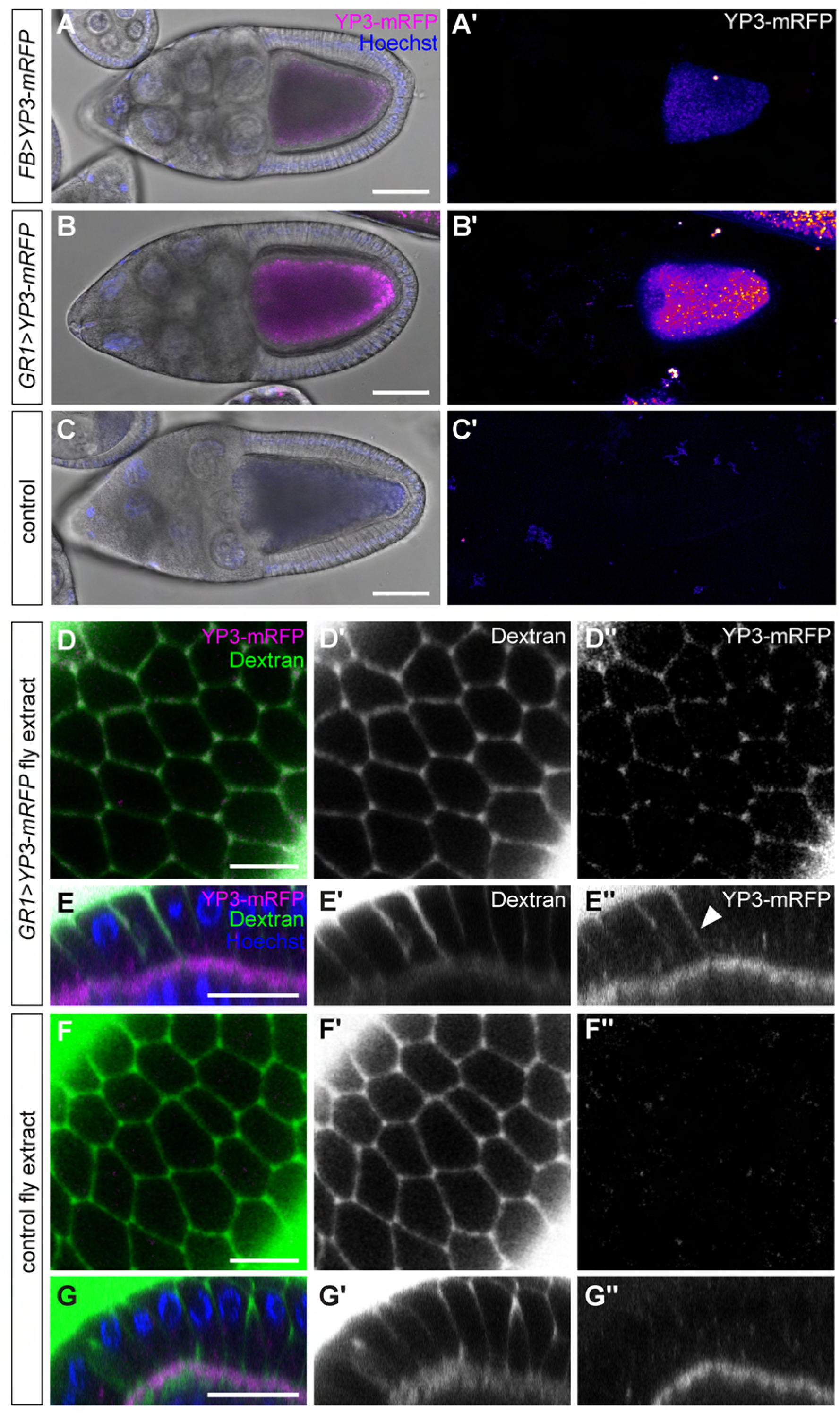
Fluorescent Yolk Protein 3 crosses the follicle epithelium in patent egg chambers. (A-C) Stage 10A egg chambers from flies expressing YP3-mRFP in the fat body (using *FB-* Gal4; A) or in follicle cells (using *GR1*-Gal4; B), or from wild-type control fly (C). Nuclei are stained with Hoechst 33342 (blue). (A-C) show transmitted light images overlaid with YP3-mRFP (magenta) and Hoechst 33342 (blue) fluorescence. (A’-B’) show Z-projections of confocal sections. Note that YP3-mRFP produced either in the fat body or in follicle cells is taken up by the oocyte and accumulates in yolk granules. (C’) shows autofluorescence of a control egg chamber not expressing YP3-mRFP. Fluorescence intensities in (A’-C’) are color-coded using the heat map shown to the right. (D-G) Stage 10A wild-type egg chambers were incubated with FITC-Dextran (40 kDa; green) and extract from flies expressing YP3-mRFP under the control of *GR1*-Gal4 (D-E) or extract from wild-type flies not expressing YP3-mRFP (F-G). En-face views (D,F) and cross-sections (E,G) are shown. Cross-sections also show nuclei stained with Hoechst 33342 (blue). Note that YP3-mRFP fills the intercellular gaps between FCs across the FCE (D”,E”), colocalizing with Dextran (D’,E’). The control egg chamber (F-G) incubated in extract from wild-type flies shows the level of autofluorescence. Scale bars: (A-C), 50 μm; (D-I), 10 μm; (D’-I’), 20 μm.

**Supplementary Figure 2.**
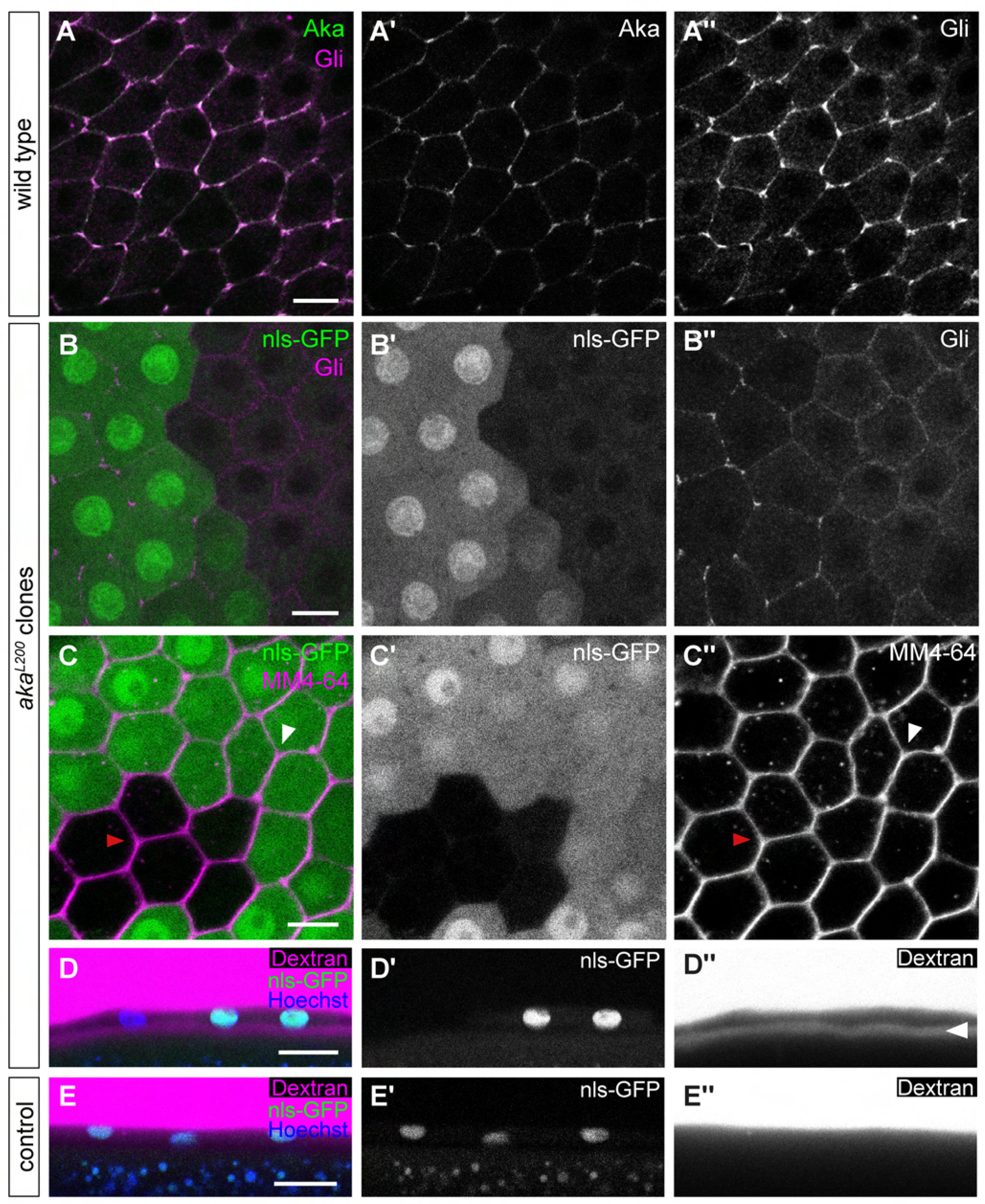
Assembly of tricellular junctions marks the termination of patency. (A) Immunostaining of wild-type late stage 10B egg chamber showing that Aka (green; A’) and Gli (magenta; A”) localize at TCJs. (B) Immunostaining of Gli (magenta) in late stage 10B mosaic egg chamber carrying an *aka^L200^* loss-of-function clone marked by the absence of nls-GFP (green). Note that Gli accumulates at TCJs in wild-type cells, but fails to accumulate at TCJs and is mislocalized along bicellular junctions in the *aka^L200^* mutant cells. (C) Confocal section of FCs in living egg chamber (late stage 10B) containing an *aka^L200^* clone marked by the absence of nls-GFP (green). Cell membranes are marked by MM4-64 (magenta). Note that FC vertices are closed in wild-type (white arrowhead) and in *aka^L2^*^00^ mutant (red arrowhead) parts of the epithelium (C’). (D-E) Cross-sections of FCE in living post-patency (stage 12) egg chambers containing *aka^L2^*^00^ clones marked by the absence of nls-GFP (green) and incubated in medium containing TIRTC-Dextran (10 kDa; magenta) and Hoechst 33342 (blue). (D-D”) shows that follicles bearing *aka^L2^*^00^ mutant clones are permeable for dextran, which is detectable in the space between the FCE and the oocyte (arrowhead in D”). (E-E”) shows a wild-type stage 12 control egg chamber from a nls-GFP FRT40A female. Note that dextran does not pass the epithelium, indicating that the FCE barrier is intact. Scale bars: (A-C), 10 μm (D-E), 20 μm.

**Supplementary Figure 3.**
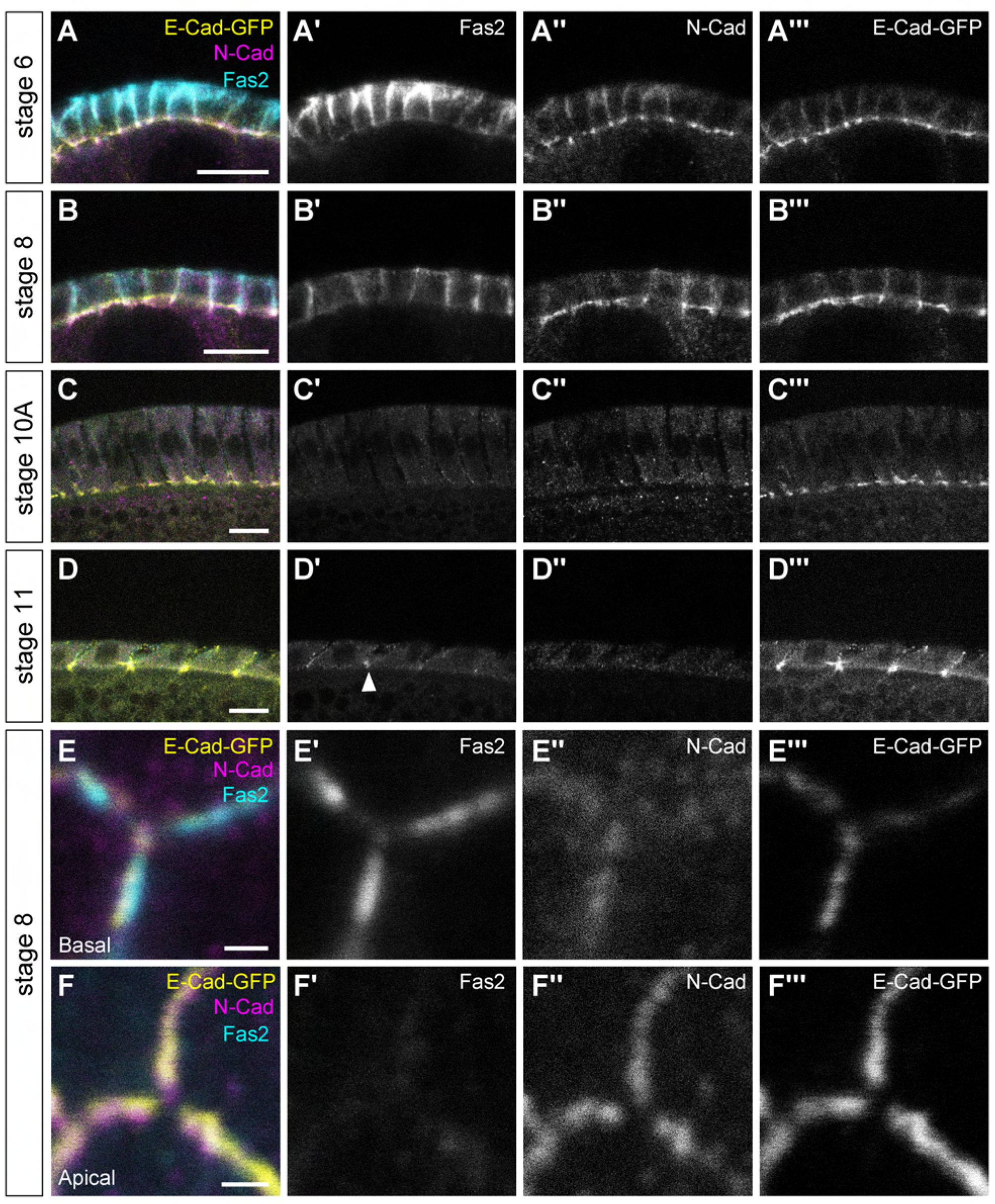
Sequential removal of adhesion proteins from vertices precedes patency. (A-F) The distribution of Fas2 (cyan; anti-Fas2 immunostaining) N-Cad (magenta; anti-N-Cad immunostaining) and E-Cad-GFPx3 (yellow; GFP fluorescence) was analyzed in fixed egg chambers expressing E-Cad-GFPx3 at pre- (stage 6; A), early (stage 8; B), mid- (stage 10A; C), and late (stage 11; D) vitellogenic stages. (A-D) show cross-sections, (E,F) show sections taken orthogonally to the apical-basal axis near the basal (E) or apical (F) surface. (A) At stage 6 Fas2 is abundant on basolateral membranes (A’), while E-Cad and N-Cad show strong accumulation at the apical adherens junctions and weaker signals along basolateral membranes (A”-A”’). (B) At stage 8 Fas2 levels are strongly reduced (B’), while E-Cad and N-Cad levels remain comparable to stage 6 (B”-B”’). (C) At stage 10A, coinciding with patency, Fas2 and N-Cad are no longer detectable at FC contacts (C’-C”), whereas E-Cad is still present at the apical side (C”). (D) At stage 11, after patency, Fas2 reappears and localizes apically (D’, arrowhead), whereas N-Cad remains absent (D”). (E,F) Sections of a FC vertex (stage 8) taken near the basal (E) or apical (F) surface. Note that basolaterally (E) Fas2 is removed selectively from vertices (E’). On the apical side both E-Cad and N-Cad are removed from vertices (F”-F”’). Scale bars: (A-D), 10 μm; (E-F), 1 μm.

## Supplementary movies

**Video S1**

**Intercellular channels in patent stage 10A follicles are permeable for dextran.**

3D animation of FCE in living stage 10A egg chamber in medium containing 40 KDa FITC-Dextran (green), Hoechst 33342 to label nuclei (blue), and MM4-64 to label cell membranes (magenta). Note that Dextran-filled channels span the epithelium from the basal side to the oocyte surface. Channels adjacent to a single follicle cell were segmented and are highlighted in grey.

**Video S2**

**Distribution of E-Cad::GFP in FCs in basolateral puncta and apical adherens junctions.**

3D animation of confocal Z-stack from a living stage 8 egg chamber expressing E-Cad::GFPx3 (green). Nuclei are labeled with Hoechst 33342 (blue). Note that E-Cad::GFPx3 accumulates strongly in a continuous belt at the apical zonula adherens (ZA) and is distributed in puncta along the basolateral membrane.

**Video S3**

**E-Cad puncta limit the open intercellular space between FCs.**

Time-lapse movie of FC vertex in a stage 9 egg chamber expressing E-Cad::GFPx3 (green). Cell membranes are labeled by MM4-64 (magenta). A confocal section at the level of the basolateral membrane is shown. Time is indicated. At t=0 seconds a hyper-osmotic shock was applied by adding PBS to the culture medium (see Materials and Methods). Membranes at the vertex separate immediately upon addition of PBS, leading to the opening of an intercellular space. Note that E-Cad::GFPx3 puncta limit the separating membranes at the boundary of the open intercellular space.

Scale bar, 1 μm.

